# KLP-6 is a kinesin superfamily protein resistant to ADP inhibition

**DOI:** 10.1101/2025.02.11.637744

**Authors:** Tomoki Kita, Shinsuke Niwa

## Abstract

Product inhibition is a type of enzyme inhibition in which the reaction product suppresses further enzyme activity. Kinesins are microtubule-dependent ATPases that move along microtubules by hydrolyzing ATP into ADP and inorganic phosphate (Pi). Like other ATPases, kinesins are inhibited by their hydrolysis product, ADP. We show here that *Caenorhabditis elegans* kinesin-3, KLP-6, is a unique kinesin that is resistant to ADP inhibition. Amino acid sequence comparisons between KLP-6 and KIF1A, another kinesin-3, revealed that KLP-6 possesses a unique sequence in an α2a helix. Substituting this region with the corresponding sequence from KIF1A abolished KLP-6’s markedly higher binding selectivity for ATP over ADP, confirming that this domain is crucial for its insensitivity to ADP inhibition. Molecular dynamics simulations uncovered that the α2a helix of KLP-6 adopts a less stable helical conformation in the ADP-bound state than in the ATP-bound state, which explains KLP-6’s strong selectivity for ATP. Our results provide valuable insights into kinesin’s nucleotide selectivity and underlying molecular mechanism.

## Introduction

Product inhibition is a common phenomenon in enzyme-catalyzed reactions (Holdgate *et al*, 2018). Since most product molecules of an enzyme reaction share structural similarities with the substrate, they can still bind to the enzyme’s active site, preventing substrate binding and reducing the reaction velocity. Product inhibition plays a crucial role in regulating cellular metabolism, serving as a negative feedback mechanism to control metabolic pathways (Sen & Schulz, 1989). Overcoming this inhibition is also important in the biotechnology industry to enhance product yield (Deu *et al*, 2012; Zhang *et al*, 2015; Fan *et al*, 2017).

Some enzymes have been reported to be resistant to product inhibition. Glucokinase, also known as hexokinase IV or D, is one of the four members of the hexokinase enzyme family. It phosphorylates glucose, using ATP as the second substrate, to form glucose-6-phosphate. Unlike other hexokinases, glucokinase has a lower affinity for glucose and is insensitive to physiological concentrations of glucose-6-phosphate (MIDDLETON, 1990). β-glucosidases hydrolyze cellooligosaccharides, such as cellobiose and cellotriose, into D-glucose during plant biomass saccharification. Some β-glucosidases are not inhibited and, surprisingly, are activated in the presence of D-glucose (Zanoelo *et al*, 2004; Uchiyama *et al*, 2013; Matsuzawa & Yaoi, 2017). Owing to their product inhibition tolerance, these β-glucosidases are considered promising candidates for improving biomass saccharification.

Most molecular motor proteins hydrolyze ATP, generating ADP and Pi, to move or rotate directionally. Myosins are well known for their roles in muscle contraction, and their activity on actin filaments is influenced by ADP product inhibition (Wang *et al*, 2000; De La Cruz *et al*, 2000). Notably, skeletal muscle myosin is less affected by ADP inhibition compared to cardiac and smooth muscle myosins, highlighting differences in how ADP modulates contractile function across muscle types (Drew *et al*, 1992; Yamashita *et al*, 1994). F1-ATPase is a rotary molecular motor in which the central γ-subunit rotates against the α3β3 cylinder. When Mg·ADP binds from the external medium, the motor enters an inhibited state and rarely releases the bound Mg·ADP (Vasilyeva *et al*, 1982; Milgrom & Boyer, 1990; Jault & Allison, 1993). A mutation in the nucleotide-binding site, known as the Walker A motif or P-loop, has been shown to reduce ADP inhibition in F1-ATPase, enabling detailed studies of its energetics (Jault *et al*, 1996; Muneyuki *et al*, 2007). In eukaryotic cilia and flagella, axonemal dyneins coordinate their functions to generate oscillatory bending of axonemes. Axonemal dynein contains four nucleotide-binding sites (Gibbons *et al*, 1991; Ogawa, 1991). Interestingly, its activity is enhanced in the presence of ADP in addition to ATP (Yagi, 2000; Shiroguchi & Toyoshima, 2001), suggesting that additional ADP binding plays a physiological role in regulating dynein activity (Ishibashi *et al*, 2020). Kinesins are primarily responsible for intracellular transport and move along microtubules. The movement of kinesin-1, also called conventional kinesin, is strongly inhibited by ADP but not by Pi (Schief *et al*, 2004). Although kinesin proteins are classified into 15 families (Hirokawa *et al*, 2009), no species exhibiting ADP inhibition tolerance, nor any mutations conferring this property, have been reported to date.

Here, we report that the *Caenorhabditis elegans* (*C. elegans*) kinesin-3, KLP-6, is insensitive to ADP inhibition and exhibits a high binding selectivity for ATP over ADP. KLP-6 is responsible for transporting a mechanosensory receptor complex in male-specific *ce*phalic *m*ale (CEM) cilia (Peden & Barr, 2005; Morsci & Barr, 2011). KLP-6 moved along microtubules at an unchanged speed in the presence of equal concentrations of ADP and ATP, in contrast to another kinesin-3 member, KIF1A, which exhibited a significantly reduced speed in the presence of ADP. Comparative analysis, as well as molecular dynamics (MD) simulations suggests that the stability of KLP-6’s unique α2a helix underlies the selectivity for ATP over ADP.

## Results

### KLP-6 exhibits directional movement along microtubules in an ADP solution

We compared the single-molecule activities of two kinesin-3 proteins, human KIF1A and *C. elegans* KLP-6, using total internal reflection fluorescence (TIRF) microscopy. To directly investigate the motility parameters of these motors, we analyzed truncated versions containing the motor and neck coiled-coil domains (Fig. 1A). Since the neck coiled-coil domains of KIF1A and KLP-6 do not form stable dimers (Okada & Hirokawa, 1999; Kita *et al*, 2024, 2025), the motors were stably dimerized via fusion to a leucine zipper (KIF1A(1-393)LZ and KLP-6(1-390)LZ) (Kita *et al*, 2024; Soppina *et al*, 2014; Budaitis *et al*, 2021; Boyle *et al*, 2021; Lam *et al*, 2021; Anazawa *et al*, 2022; Kita *et al*, 2023) (Fig. 1A).

**Figure 1.**
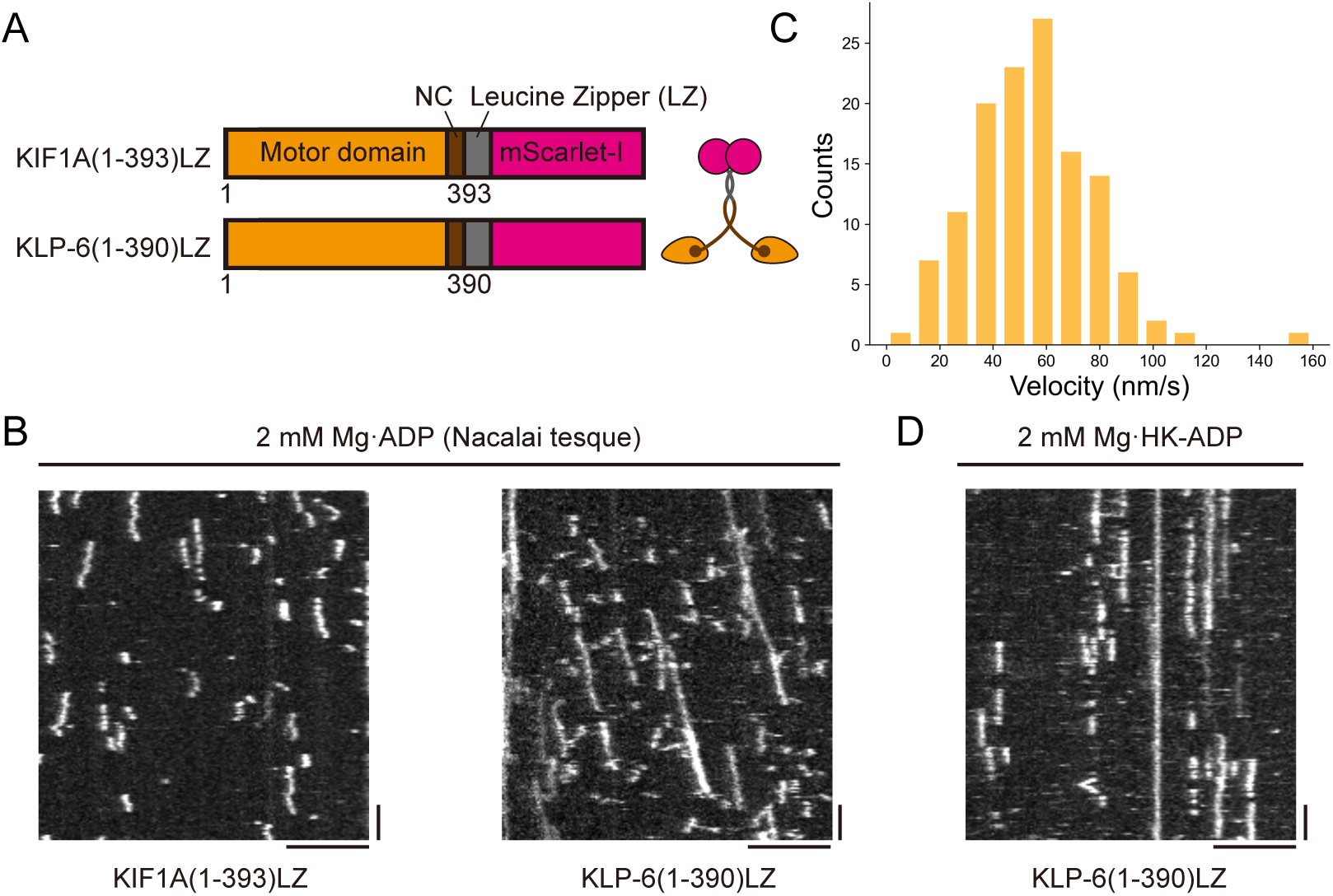
Directional movements of KLP-6(1-390)LZ in an ADP solution. (A) Schematic illustration of the domain organization in KIF1A(1-393)LZ and KLP-6(1-390)LZ tagged with a fluorescent protein mScarlet-I. (B) Representative kymographs showing the motility of KIF1A(1-393)LZ and KLP-6(1-390)LZ in the presence of 2 mM Mg·ADP purchased from Nacalai tesque. Scale bars: horizontal, 5 µm; vertical, 10 seconds. (C) Histogram of velocity for KLP-6(1-390)LZ in the presence of 2 mM Mg·ADP. n = 129 (D) Representative kymograph showing the motility of KLP-6(1-390)LZ in the presence of 2 mM Mg·HK-ADP. Scale bars: horizontal, 5 µm; vertical, 10 seconds.

Normally, kinesins lack processivity along microtubules in ADP solution (Lu *et al*, 2009). A well-studied KIF1A(1-393)LZ did not exhibit unidirectional movement in the presence of 2 mM Mg·ADP purchased from Nacalai Tesque (Anazawa *et al*, 2022; Kita *et al*, 2023) (Fig. 1B). Surprisingly, we observed that KLP-6(1-390)LZ moved along microtubules at a speed of 55±2 nm/s (mean±S.E.) in the same condition (Figs. 1B and C). Although the speed was much slower than in the presence of 2 mM Mg·ATP (260 nm/s) (Kita *et al*, 2024), KLP-6 obviously possessed processivity even in ADP solutions. One possible explanation for this movement is that KLP-6(1-390)LZ efficiently hydrolyzes trace amounts of ATP that was originally present as a contaminant in the purchased ADP stock (Hancock & Howard, 1999; Motojima & Yoshida, 2003). To test this hypothesis, we depleted ATP from the ADP stock using hexokinase (You *et al*, 2017) and observed the motility of KLP-6(1-390)LZ in a hexokinase-treated ADP (HK-ADP) solution. As a result, KLP-6(1-390)LZ lost its processivity (Fig. 1D), suggesting that KLP-6 can bind trace amounts of ATP even in the presence of excess ADP.

### KLP-6 is minimally affected by ADP inhibition

To quantify ADP inhibition in KLP-6, we measured motor velocity on microtubules at various Mg·ATP concentrations ([Mg·ATP]) in the presence or absence of 1 mM Mg·HK-ADP. Under all conditions used in this study, the dependence of the single-molecule velocity on [Mg·ATP] was well described by the standard Michaelis–Menten equation (Schief *et al*, 2004):

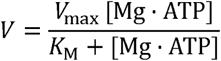

where *V*_max_ represents the maximum speed, and *K*_M_ denotes the [Mg·ATP] concentration required to achieve half-maximal velocity. In the presence of Mg·HK-ADP, *V* is described as

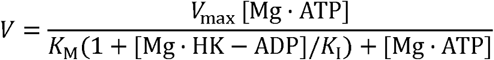

where *K*_I_ represents the inhibition constant for Mg·HK-ADP.

First, we analyzed KIF1A(1-393)LZ as a control. In the presence of 1 mM ADP and ATP, the motor velocity was significantly reduced compared to that observed in the ATP-only solution (Figs. 2A and B), consistent with previous observations for the conventional kinesin (Schief *et al*, 2004). In contrast, the velocity of KLP-6(1-393)LZ remained unchanged, even in the presence of 1 mM ADP and ATP (Figs. 2C and D). From these plotting, we determined the kinetic parameters for KIF1A(1-393)LZ: *V*_max_ = 2381±23 nm/s (mean±S.E.), *K*_M_ = 190±3 µM (mean±S.E.), and *K*_I_ = 247±3 µM (mean±S.E.) (Figs. 2E-G and Table. 1). *V*_max_ and *K*_M_ are in agreement with prior studies (Soppina *et al*, 2014; Budaitis *et al*, 2021; Boyle *et al*, 2021; Zaniewski & Hancock, 2023). For KLP-6(1-390)LZ, the corresponding parameters were *V*_max_ = 248±3 nm/s (mean±S.E.) and *K*_M_ = 20.1±0.5 µM (mean±S.E.) which are comparable to those of *C. elegans* kinesin-2 (Pan *et al*, 2006) and conventional kinesin (Schief *et al*, 2004), respectively (Figs. 2E, F and Table. 1). In contrast, the *K*_I_ of KLP-6(1-390)LZ was 901±31 µM (mean±S.E.) (Fig. 2G), the highest value reported among the kinesin superfamily proteins measured to date. We also measured *K*_I_ in the presence of different ADP concentrations (2 mM and 5 mM). The values were largely consistent: *K*_I_ values in the presence of 2 mM and 5 mM ADP for KIF1A(1-393)LZ were 180 ± 4 µM and 214 ± 6 µM, and for KLP-6(1-390)LZ were 1000 ± 69 µM and 1051 ± 44. (mean±S.E.) (Fig. S1). We therefore used the *K*_I_ value measured in the presence of 1 mM ADP as the representative value throughout this article.

**Figure 2.**
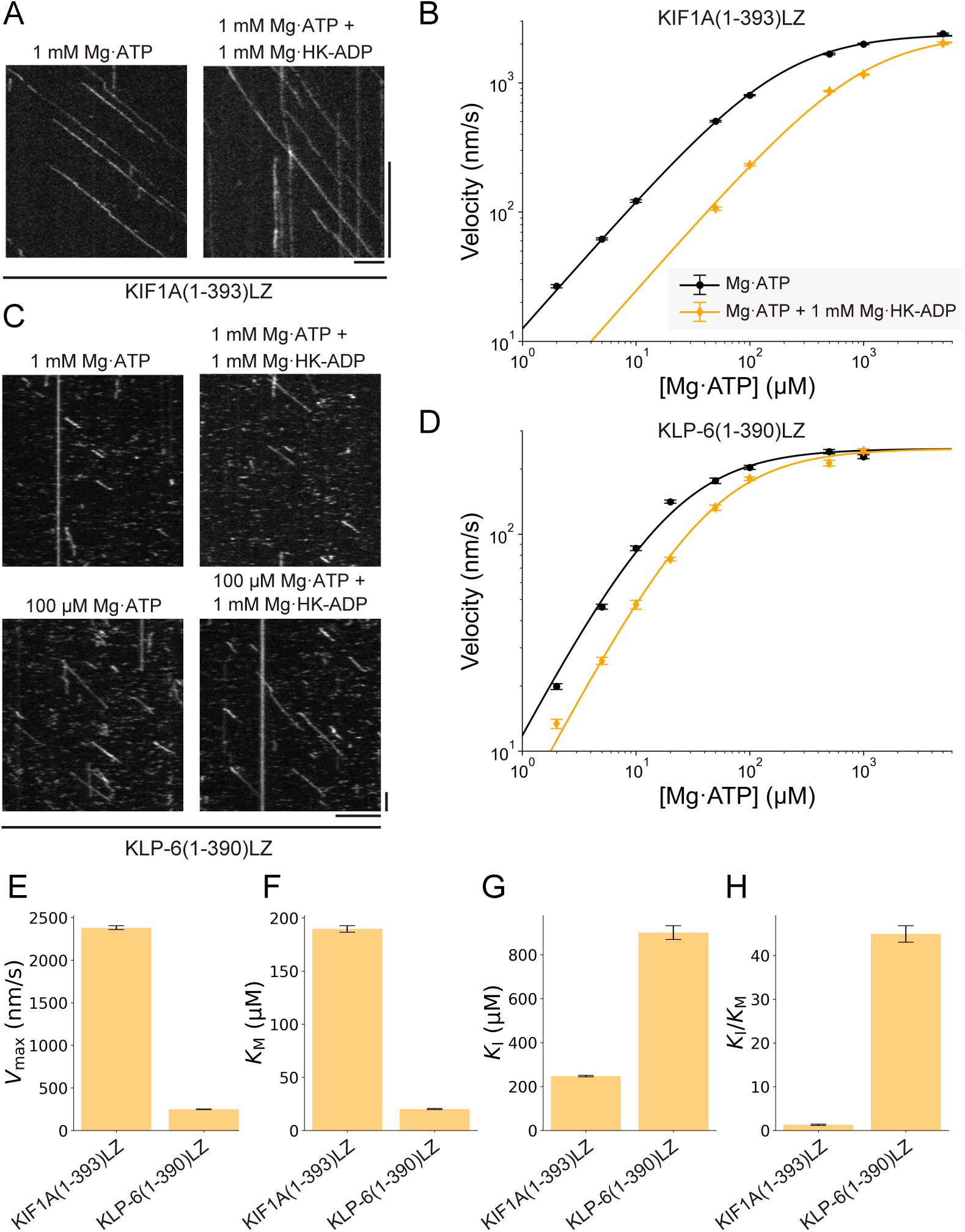
Minimum ADP inhibition in KLP-6(1-390)LZ. (A) Representative kymographs showing the motility of KIF1A(1-393)LZ in the presence of 1 mM Mg·ATP, with or without an additional 1 mM Mg·HK-ADP. Scale bars: horizontal, 5 µm; vertical, 10 seconds. (B) Single-molecule velocity of KIF1A(1-393)LZ versus [Mg·ATP] in the absence (circles) and presence (diamonds) of 1 mM HK-ADP. The black and orange solid lines represent the regression curves fitted to the Michaelis-Menten equation and the equation with product inhibition, respectively. Each data point represents the mean ± S.E. At least three independent experiments were conducted for each condition, with more than 114 motile particles analyzed. (C) Representative kymographs showing the motility of KLP-6(1-390)LZ in the presence of 1 mM Mg·ATP, with or without an additional 1 mM Mg·HK-ADP, and in the presence of 100 μM Mg·ATP, with or without an additional 1 mM Mg·HK-ADP. Scale bars: horizontal, 5 µm; vertical, 10 seconds. (D) Single-molecule velocity of KLP-6(1-390)LZ versus [Mg·ATP] in the absence (circles) and presence (diamonds) of 1 mM HK-ADP. The black and orange solid lines represent the regression curves fitted to the Michaelis-Menten equation and the equation with product inhibition, respectively. Each data point represents the mean ± S.E. At least three independent experiments were conducted for each condition, with more than 131 motile particles analyzed. (E-H) Comparison of (E) *V*_max_, (F) *K*_M_ (G) *K*_I_ and (H) *K*_I_*/K*_M_ between KIF1A(1-393)LZ and KLP-6(1-390)LZ. Error bars represent the mean ± S.E.

**Table 1.** Summary of the kinetic parameters.

|  | CeKinesin-3 |  |  |  | HsKinesin-3 | DmKinesin-1* | CeKinesin-2# |
| --- | --- | --- | --- | --- | --- | --- | --- |
|  | KLP-6 | KLP-6<br>(L3SW) | KLP-6<br>(F120K) | KLP-6<br>(SIV/TM<br>M) | KIF1A | KIF5B | KLP-11/KLP-20/KAP-1 |
| $V_{\max}$<br>(nm/s) | 248±3 | 203±5 | 338±4 | 234±3 | 2381±23 | 906±9 | 260 |
| $K_M$ (μM) | 20.1±0.5 | 31.3±1.7 | 20.3±0.5 | 4.4±0.1 | 190±3 | 28.1±0.9 | 280 |
| $K_I$ (μM) | 901±31<sup>\$</sup> | 1175±78<sup>\$</sup> | 242±4<sup>\$</sup> | 11.9±0.3<sup>\$</sup> | 247±3<sup>\$</sup> | 34.6±2.6 | 42 |
| $K_I/K_M$ | 44.9±1.9<sup>\$</sup> | 37.5±2.9<sup>\$</sup> | 11.9±0.6<sup>\$</sup> | 2.7±0.3<sup>\$</sup> | 1.3±0.1<sup>\$</sup> | 1.2±0.1 | 0.15 |

Values are mean±S.E.
\*, data from (Schief *et al*, 2004)
#, data from (Pan *et al*, 2006)

To quantify the binding selectivity for ATP over ADP, we defined the binding selectivity ratio as *K*_I_ */K*_M_. Although this ratio does not directly correspond to the binding constant ratio of the ADP and ATP (*K*_ADP_*/K*_ATP_), it serves as a useful proxy (Drew *et al*, 1992). The selectivity ratio for KIF1A was 1.3±0.1 (mean±S.E.), which is comparable to that of conventional kinesin (Schief *et al*, 2004) (Fig. 2H and Table. 1). This indicates that KIF1A is significantly affected by ADP inhibition and exhibit no strong preference between ATP and ADP binding. In contrast, due to the high *K*_I_ value of KLP-6(1-390)LZ, the selectivity ratio of KLP-6(1-390)LZ was 44.9±1.9 (mean±S.E.), which is much higher than that of KIF1A(1-393)LZ (Fig. 2H and Tabel. 1). These findings highlight that KLP-6 is a unique kinesin that exhibits remarkable resistance to ADP inhibition.

### Unique amino acid sequence of KLP-6

To identify the regions responsible for ADP inhibition tolerance in KLP-6, we compared its amino acid sequence with those of other kinesin-3s from humans and *C. elegans* (Fig. 3A). A notable distinction of KLP-6 compared to other kinesin-3s is a unique insertion in loop-3. Unlike the loop-3 of KIF1A, the longer loop-3 of KLP-6 extends over the nucleotide-binding site (Kikkawa *et al*, 2001; Wang *et al*, 2022) (Figs. 3B and C). Kinesin-3 family proteins naturally possess an extended loop-3 (Vale, 2003), and the additional extension of this loop in KLP-6 may contribute to its significantly higher binding preference for ATP over ADP.

**Figure 3.**
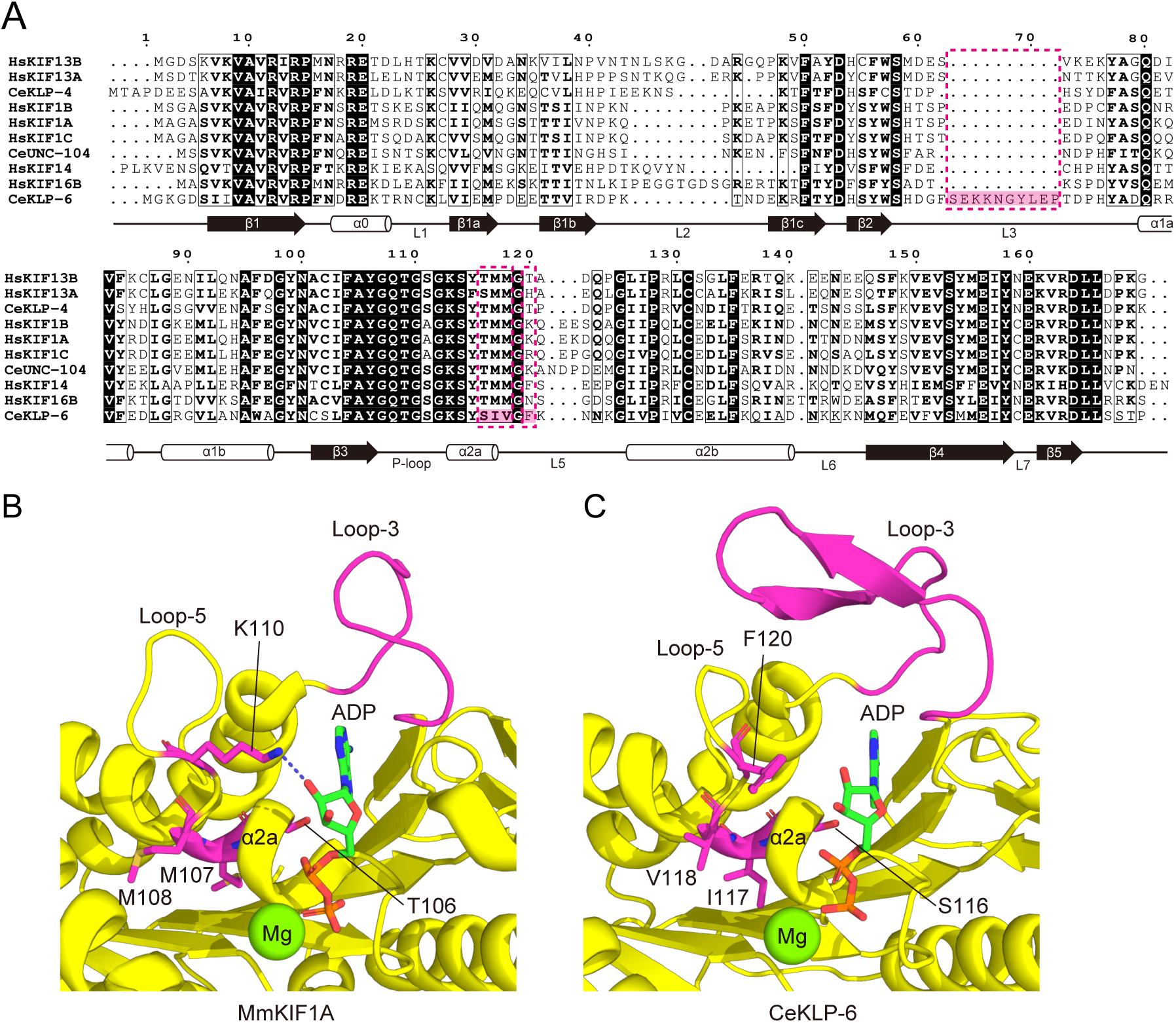
Amino acid sequence and structural comparison between KLP-6 and other kinesin-3s. (A) Alignment of kinesin-3 amino acid sequences. Identical residues are highlighted in black, while highly conserved residues are outlined with a black border. The secondary structures of KLP-6 are indicated at the bottom. KLP-6-specific residues are highlighted in magenta. (B) Protein structure of the motor domain of *Mus musculus* KIF1A bound to Mg·ADP (PDB ID 1I5S). Loop-3 and highly conserved residues (T106, M107, and M108) are highlighted in magenta. K110, also highlighted in magenta, forms a hydrogen bond with the ADP-ribose. (C) Protein structure of the motor domain of *C. elegans* KLP-6 bound to Mg·ADP (PDB ID 7WRG). This structure was derived from the full-length KLP-6 structure, with all domains other than the motor domain (with Mg·ADP) removed for visualization. Loop-3 and KLP-6-specific residues (S116, I117, and V118) are highlighted in magenta. F120, corresponding to K110 in KIF1A, is also highlighted in magenta.

The P-loop contains the consensus sequence G*X*_4_GK(T/S). In F1-ATPase, replacing the C-terminal threonine in the P-loop of the β subunit with serine has been shown to reduce ADP inhibition (Jault *et al*, 1996; Muneyuki *et al*, 2007). In kinesin-3 family members, the P-loop naturally terminates with serine; however, differences in the adjacent α2a helix distinguish KLP-6 from other kinesin-3 proteins (Fig. 3A). In most kinesin-3s, the conserved sequence at the end of the α2a helix is TMM, whereas KLP-6 uniquely features the sequence SIV (Fig. 3A). The first threonine and second methionine are also conserved in kinesin-1 and kinesin-2 family members (Fig. S2), suggesting that this sequence plays a crucial role in nucleotide binding and/or ATP hydrolysis.

From a structural perspective, the loop-5 of KIF1A contains a lysine (K110) that forms a hydrogen bond with ADP-ribose, the corresponding position in KLP-6 is occupied by phenylalanine (F120), which lacks this interaction (Figs. 3B and C). This destabilization of nucleotide binding may contribute to KLP-6’s high resistance to ADP inhibition.

### Unique sequence in the **α**2a helix contribute to minimum sensitivity to ADP inhibition in KLP-6

Based on sequence comparisons between KLP-6 and other kinesin-3 proteins, we generated three mutant KLP-6 constructs (Figs. 4A and B) and analyzed the impact of ADP inhibition on their activity (Figs. 4C-F). First, we tested whether the long loop-3 is responsible for the KLP-6’ s high binding selectivity for ATP over ADP. A ‘swap’ construct, KLP-6(1-390)LZ(L3SW), was created by replacing the native loop-3 of KLP-6 with the loop-3 from KIF1A (Figs. 4A and B). While the velocity in the presence of 1 mM Mg·ATP was unaffected by the addition of 1 mM HK-Mg·ADP, similar to the wild type (Figs. 4C and D), the loop-3 swap caused a slight decrease in *V*_max_ to 203±5 nm/s (mean±S.E.) and increases in both *K*_M_ and *K*_I_, measured at 31.3±1.7 µM and 1175±78 µM (mean±S.E.), respectively (Figs. 4G-I and Table. 1). Consequently, the binding selectivity ratio (*K*_I_ */K*_M_) was calculated to be 37.5±2.9 (mean±S.E.), representing a slight decrease compared to the wild type (Fig. 4J and Table. 1). These results suggest that the extended loop-3 affects the nucleotide affinity. However, since KLP-6(1-390)LZ(L3SW) still exhibited significantly higher binding preference for ATP over ADP (Fig. 4J), loop-3 is not the primary determinant of KLP-6’s ADP inhibition tolerance.

**Figure 4.**
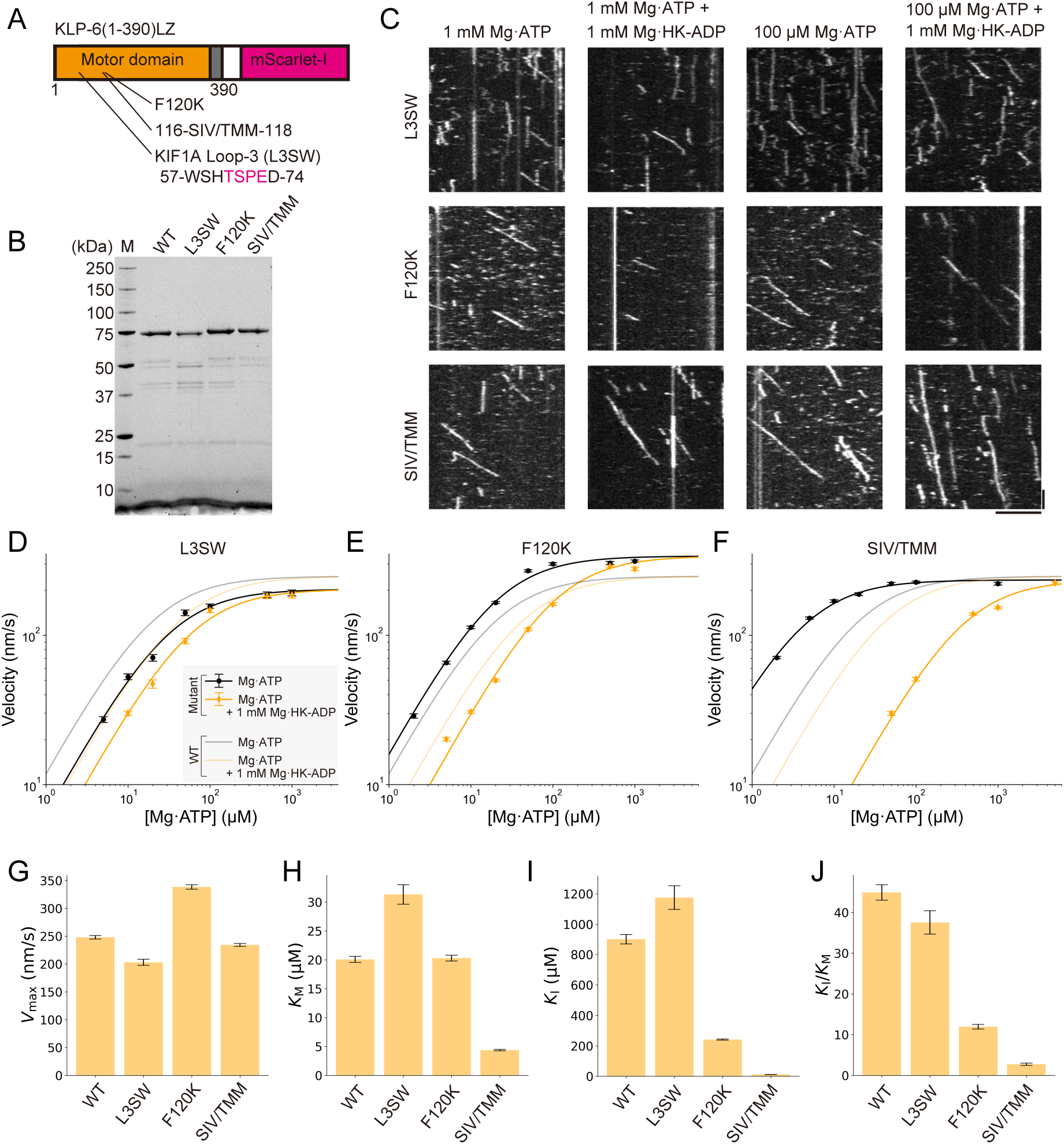
Role of domains in the minimal ADP inhibition of KLP-6(1-390)LZ. (A) Mutation sites in KLP-6(1-390)LZ. In KLP-6(1-390)LZ(L3SW), amino acids 60–73 are replaced with amino acids 60–63 from KIF1A. (B) SDS-PAGE analysis of KLP-6(1-390)LZ and its mutants. M represents the marker lane. Numbers on the left indicate molecular weight (kDa). (C) Representative kymographs showing the motility of KLP-6(1-390)LZ(L3SW), KLP-6(1-390)LZ(F120K), and KLP-6(1-390)LZ(SIV/TMM) in the presence of 1 mM Mg·ATP, with or without an additional 1 mM Mg·HK-ADP, and in the presence of 100 μM Mg·ATP, with or without an additional 1 mM Mg·HK-ADP. Scale bars: horizontal, 5 µm; vertical, 10 seconds. (D-F) Single-molecule velocity of KLP-6(1-390)LZ(L3SW), KLP-6(1-390)LZ(F120K), and KLP-6(1-390)LZ(SIV/TMM) versus [Mg·ATP] in the absence (circles) and presence (diamonds) of 1 mM HK-ADP. The black and orange solid lines represent the regression curves fitted to the Michaelis-Menten equation and the equation with product inhibition, respectively. The faint black and orange solid lines indicate the results for KLP-6(1-390)LZ, as shown in Fig. 2D. Each data point represents the mean ± S.E. At least three independent experiments were conducted for each condition, with more than 150 motile particles analyzed." (G-J) Comparison of (E) *V*_max_, (F) *K*_M_ (G) *K*_I_ and (H) *K*_I_*/K*_M_ between KLP-6(1-390)LZ, KLP-6(1-390)LZ(L3SW), KLP-6(1-390)LZ(F120K), and KLP-6(1-390)LZ(SIV/TMM). Error bars represent the mean ± S.E.

Next, we introduced the F120K mutation into KLP-6(1-390)LZ (Figs. 4A and B), which was expected to create a hydrogen bond with ADP (Fig. 3B). Interestingly, F120K turned out to be a gain-of-function mutation, increasing *V*_max_ to 338±4 nm/s (mean±S.E.) (Figs. 4C, E, G, and Table. 1). In the presence of 1 mM ADP and ATP, KLP-6(1-390)LZ(F120K) was not inhibited by ADP, consistent with the wild type (Figs. 4C and E). However, in the presence of 1 mM ADP and 100 µM ATP, the reduction in motor velocity was more pronounced compared to the wild type (Figs. 4C and E). Further analysis revealed that while its *K*_M_ of 20.3±0.5 µM (mean±S.E.) was comparable to that of the wild type (Fig. 4H and Table. 1), its *K*_I_ was significantly reduced to 242±4 µM (mean±S.E.), resulting in decreased selectivity for ATP over ADP (*K*_I_*/K*_M_ = 11.9±0.6, mean±S.E.) (Figs. 4I, J, and Table. 1). These findings indicate that the loop-5 of KLP-6 is designed to avoid forming a hydrogen bond with ADP, contributing to its resistance to ADP inhibition.

Finally, we investigated the role of KLP-6’s unique α2a helix sequence (SIV motif) in ADP inhibition resistance. Three mutations, S116T, I117M, and V118M, were introduced into KLP-6(1-390)LZ, generating the construct KLP-6(1-390)LZ(SIV/TMM) (Figs. 4A and B). In the presence of 1 mM ADP and ATP, we observed that KLP-6(1-390)LZ(SIV/TMM) was inhibited by ADP, similar to KIF1A (Figs. 4C and F). Notably, the *K*_M_ decreased approximately fivefold to 4.4±0.1 µM (mean±S.E.), and the *K*_I_ was significantly reduced by a factor of 76 to 11.9±0.3 µM (mean±S.E.), while the *V*_max_ remained largely unchanged at 234±3 nm/s (mean±S.E.) (Figs. 4G-I and Table. 1). The *K*_I_ */K*_M_ ratio was calculated to be 2.7±0.3 (mean±S.E.), which is largely consistent with KIF1A and conventional kinesin (Fig. 4J and Table. 1). These results demonstrate that the SIV motif within the α2a helix of KLP-6 is a key determinant of its higher binding preference for ATP over ADP.

### An unstable **α**2a helix weakens ADP binding in the KLP-6 protein

To elucidate the mechanism underlying the resistance to ADP inhibition observed in KLP-6, we performed MD simulations. Although MD simulations ideally require structures of KLP-6 bound to ADP or ATP in complex with a microtubule, only the autoinhibited ADP-bound structure is currently available (Wang *et al*, 2022) (Fig. S3A). Given the high structural conservation between the motor domains of KLP-6 and KIF1A (Fig. S3B), we employed homology modeling to generate microtubule-bound KLP-6 structures in each nucleotide state (Šali & Blundell, 1993) (Figs. S3C and D), based on the cryo-EM structures of KIF1A (ATP- and microtubule-bound: PDB ID 8UTN; ADP- and microtubule-bound: PDB ID 8UTR) (Benoit *et al*, 2024). For each nucleotide state, ten independent simulations of 100 ns were performed, and the root mean square fluctuation (RMSF) of the nucleotide was calculated to assess binding stability.

In the ATP-bound state, ATP remained stably associated with the P-loop throughout the 100 ns simulation (Fig. 5A and Movie 1). In contrast, in the ADP-bound state, ADP exhibited greater fluctuations (Figs. 5B and Movie 2). We investigated the relationship between nucleotide binding stability and the SIV motif within the α2a helix. Since the SIV motif did not form hydrogen bonds with either ATP or ADP (Fig. S4), these residues were not directly involved in nucleotide binding. Notably, the solvent environment surrounding the SIV motif appeared less favorable to the motif in the ADP-bound state than in the ATP-bound state. Since the nucleotide binding pocket adopted an open conformation in the ADP-bound state (Benoit *et al*, 2024), the α2a helix shifts closer to the β1 and β8 sheets and farther from the β6 and β7 sheets (Figs. S5A-D). This conformation resulted in reduced solvent exposure of the hydrophilic residue S116 and increased exposure of the hydrophobic residue I117 within the SIV motif (Fig. S5E). Under these conditions, secondary structure analysis revealed that the α2a helix was more frequently disrupted in the ADP-bound state than in the ATP-bound state (Fig. S6, Movies 1 and 2). The probability of the α2a helix maintaining its helical conformation over time was inversely correlated with ADP fluctuation (Fig. 5C).

**Figure 5.**
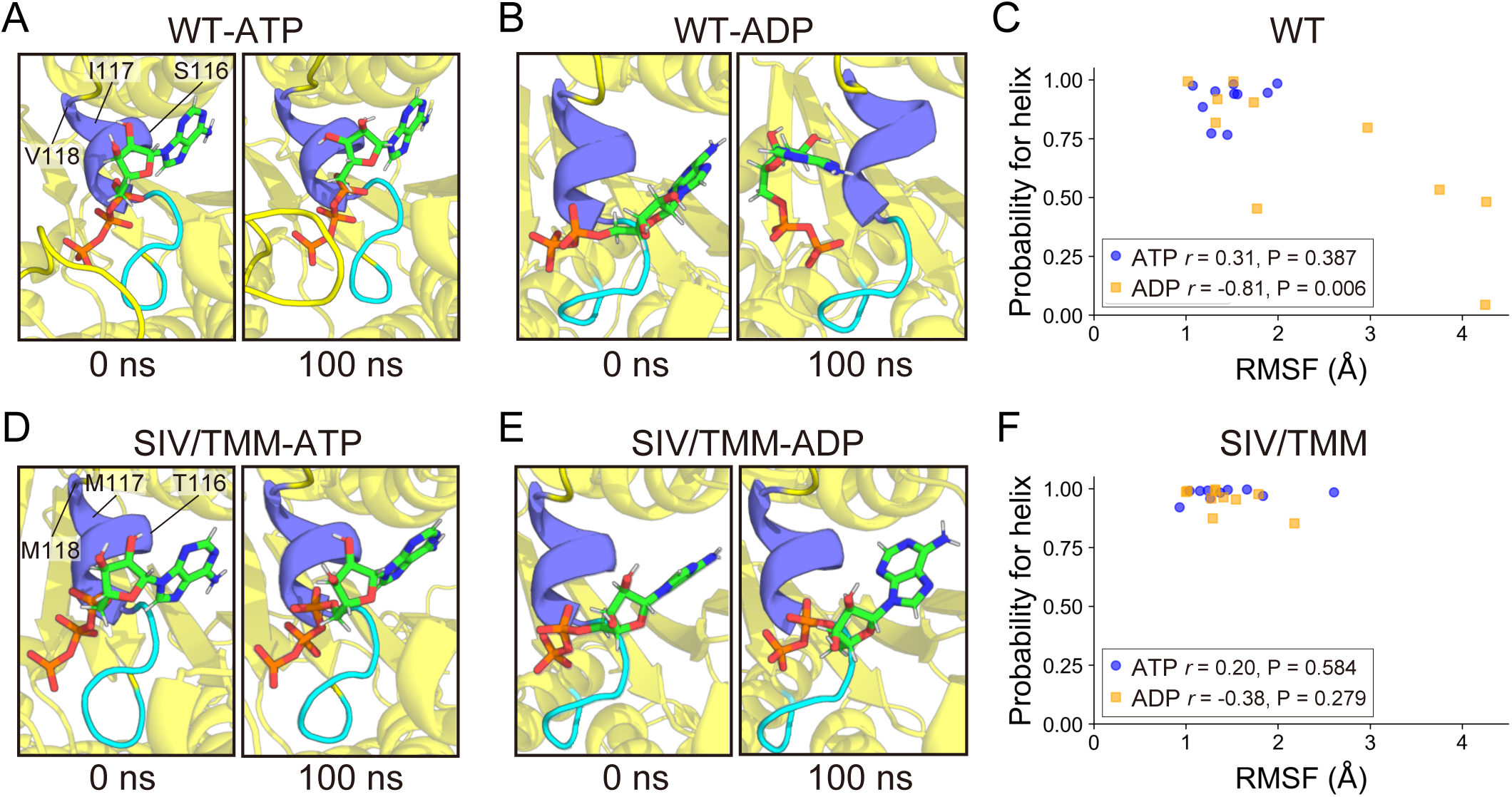
Mechanism underlying resistance to ADP inhibition in KLP-6 revealed by MD simulations. (A and B) Representative structures at 0 ns and 100 ns from MD simulations of the ATP-bound (A) and ADP-bound (B) states of KLP-6. The P-loop and α2a helix are highlighted in light blue and indigo, respectively. (C) Scatter plots showing the relationship between nucleotide RMSF and the probability of the α2a helix maintaining its helical conformation in KLP-6. Each dot represents an individual simulation. Blue circles and orange squares represent results for the ATP-bound and ADP-bound states of KLP-6, respectively. Spearman’s correlation coefficient and P-value are indicated. (D and E) Representative structures at 0 ns and 100 ns from MD simulations of the ATP-bound (D) and ADP-bound (E) states of KLP-6(SIV/TMM). The color scheme is the same as in panels (A) and (B). (F) Scatter plots showing the relationship between nucleotide RMSF and the probability of the α2a helix maintaining its helical conformation in KLP-6(SIV/TMM). Each dot represents an individual simulation. Blue circles and orange squares represent results for the ATP-bound and ADP-bound states of KLP-6(SIV/TMM), respectively. Spearman’s correlation coefficient and P-value are indicated.

Isoleucine and valine are β-branched amino acids that preferentially form β-sheets rather than α-helices (Chou & Fasman, 1979). In contrast, methionine has a strong propensity to form α-helices, and threonine also favors α-helix formation more than serine (Chou & Fasman, 1979). Based on these properties, the SIV/TMM mutations were expected to restore the disrupted α2a helix in KLP-6. As anticipated, MD simulations revealed enhanced structural stability of the α2a helix in both ATP- and ADP-bound mutant structures (Figs. 5D-F, S6, Movies 3 and 4). Furthermore, these mutations suppressed ADP fluctuations, and no notable differences in the molecular behavior of ATP and ADP were observed (Figs. 5D-F, Movies 3 and 4), consistent with experimental findings (Fig. 4J). These results suggest that the α2a helix of KLP-6 adopts a less stable conformation in the ADP-bound state compared to the ATP-bound state, contributing to its higher binding preference for ATP over ADP.

## Discussion

In this study, we found that KLP-6 is resistant to ADP inhibition and uncovered the molecular mechanism underlying this unique characteristic. Single-molecule experiments identified the SIV motif in the α2a helix as a key determinant of KLP-6’s minimal sensitivity to ADP inhibition (Fig. 4J). MD simulations further revealed that this motif destabilizes the α2a helix more in the ADP-bound state than in the ATP-bound state (Fig. 5C). These findings suggest that structural stability near the substrate-binding site is a critical factor contributing to reduced sensitivity to product inhibition. Similarly, Han *et al*. have demonstrated that introducing a mutation that increased the flexibility of flaps near the substrate-binding site in chorismate-pyruvate lyase reduced affinity for both the substrate and product, but successfully alleviated product inhibition (Han *et al*, 2016). Together, these results provide valuable insights into the mechanisms of product inhibition tolerance in enzymes.

We additionally identified other domains in KLP-6 that influence its nucleotide affinity. Loop-5 also contributes to tolerance against ADP inhibition (Fig. 4J). While a lysine residue in KIF1A’s loop-5 forms a hydrogen bond with ADP, this interaction is absent in KLP-6 (Fig. 3). Although the KLP-6(F120K) mutation in loop-5 increased affinity for ADP (Fig. 4I), ATP affinity remained largely unchanged (Fig. 4H). MD simulations of the wild-type structures revealed that ATP is already stabilized by more hydrogen bonds than ADP due to its additional phosphate group (γ-Pi) (Fig. S4), suggesting that the extra hydrogen bond introduced by F120K has a less effect on ATP binding than ADP binding.

MD simulations also provided insights into the role of KLP-6’s extended loop-3. Single-molecule experiments showed that deletion of this loop caused a slight decrease in affinity for both ATP and ADP (Figs. 4H ans I). Our simulations indicated that although this loop does not form significant hydrogen bonds with the nucleotide (Fig. S4), it creates a stable covering pocket over the binding site that rarely adopts an open conformation (Fig. S7). These results suggest that the extended loop-3 regulates nucleotide dynamics, such as facilitating nucleotide rebinding after detachment, rather than directly influencing nucleotide affinity at the binding site. Notably, this loop does not appear to have specificity for either ATP or ADP, as its contribution to ADP inhibition tolerance was minimal (Fig. 4J).

Since the ATP/ADP ratio in cells is typically around 10 (Traut, 1994), most kinesin motors are expected to be largely unaffected by ADP inhibition in vivo. However, the ATP/ADP ratio in CEM cilia remains unknown, and the significance of KLP-6’s insensitivity to ADP inhibition is unclear. Unlike KLP-6, *C. elegans* kinesin-2, a ciliary kinesin, is highly susceptible to ADP inhibition (Pan *et al*, 2006) (Table. 1), suggesting that minimal sensitivity to ADP inhibition is not a common feature among ciliary kinesins. KLP-6 functions as a regulator of OSM-3 kinesin’s velocity and transport (Morsci & Barr, 2011); thus, maintaining a stable transport velocity, even in the presence of local ADP accumulation in the nonequilibrium cellular environment, may be crucial for efficient transport in CEM cilia.

Finally, we believe that studying KLP-6 will offer deeper insights into ATP recognition and its coupling to motor function in kinesins. KLP-6 may also serve as a valuable model for developing molecular motors resistant to ADP inhibition, potentially advancing research on motor energetics (Muneyuki *et al*, 2007) and contributing to the development of efficient molecular motor–based robots and systems (Akter *et al*, 2022; Kawamata *et al*, 2024).

## Methods

### Expression of KLP-6 proteins

Reagents were purchased from Nacalai tesque (Kyoto, Japan), unless described. Expression of KLP-6 proteins was performed as described (Kita *et al*, 2024). The mutants were generated through PCR-based mutagenesis using KOD plus neo DNA polymerase (TOYOBO, Tokyo, Japan). To express KLP-6 (1–390)::LZ::mScarlet-I::Strep-tag II and its mutants, LOBSTR(DE3) was transformed and selected on an LB agar plate supplemented with kanamycin at 37°C overnight. Colonies were picked and cultured in 10 mL LB medium supplemented with kanamycin overnight. Next morning, 5 mL of the medium was transferred to 500 mL 2.5×YT (20 g/L tryptone, 12.5 g/L yeast extract, 6.5 g/L NaCl) supplemented with 10 mM phosphate buffer (pH 7.4) and 50 μg/mL kanamycin in a 2 L flask and shaken at 37°C. When OD600 reached 0.6, flasks were cooled in ice-cold water for 30 min. Then, 23.8 mg IPTG was added to each flask. Final concentration of IPTG was 0.2 mM. Flasks were shaken at 18°C overnight. Next day, bacteria expressing recombinant proteins were pelleted by centrifugation (3000×g, 10 min, 4°C), resuspended in PBS and centrifuged again (3000×g, 10 min, 4°C). Pellets were resuspended in protein buffer (50 mM HEPES-KOH, pH 8.0, 150 mM KCH3COO, 2 mM MgSO4, 1 mM EGTA, 10% glycerol) supplemented with phenylmethylsulfonyl fluoride (PMSF).

### Purification of KLP-6 proteins

Purification of KLP-6 proteins was performed as described (Kita *et al*, 2024). Bacteria were lysed using a French Press G-M (Glen Mills, NJ, USA) as described by the manufacturer. After being incubated with 1% streptomycin sulfate on ice for 20 min to eliminate nucleic acids from protein samples, lysates were cleared by centrifugation (75,000×g, 20 min, 4°C) and subjected to affinity chromatography described below. Lysate was loaded on Streptactin-XT resin (IBA Lifesciences, Göttingen, Germany) (bead volume: 2 mL). The resin was washed with 40 mL Strep wash buffer (50 mM HEPES-KOH, pH 8.0, 450 mM KCH3COO, 2 mM MgSO4, 1 mM EGTA, 10% glycerol). Protein was eluted with 40 mL Strep elution buffer (50 mM HEPES-KOH, pH 8.0, 150 mM KCH3COO, 2 mM MgSO4, 1 mM EGTA, 10% glycerol, 300 mM biotin). Eluted solution was concentrated using an Amicon Ultra 15 (Merck, Darmstadt, Germany) and then separated on an NGC chromatography system (Bio-Rad, Hercules, CA, USA) equipped with a Superdex 200 Increase 10/300 GL column (Cytiva, Tokyo, Japan). Peak fractions were collected and concentrated using an Amicon Ultra 4 (Merck). Proteins were analyzed by SDS-PAGE. Concentrated proteins were aliquoted and snap-frozen in liquid nitrogen.

### Preparation of microtubules

Tubulin was purified from porcine brain as described (Castoldi & Popov, 2003). Tubulin was labeled with Biotin-PEG_2_-NHS ester (Tokyo Chemical Industry, Tokyo, Japan) and AZDye647 NHS ester (Fluoroprobes, Scottsdale, AZ, USA) as described (Kita & Niwa, 2024). To polymerize Taxol-stabilized microtubules labeled with biotin and AZDye647, 30 μM unlabeled tubulin, 1.5 μM biotin-labeled tubulin and 1.5 μM AZDye647-labeled tubulin were mixed in BRB80 buffer (80 mM PIPES, 1mM MgCl_2_, and 1mM EGTA) supplemented with 1 mM GTP and incubated for 15 min at 37°C. Then, an equal amount of BRB80 supplemented with 40 μM taxol was added and further incubated for more than 15 min. The solution was loaded on BRB80 supplemented with 300 mM sucrose and 20 μM taxol and ultracentrifuged at 100,000 g for 5 min at 30°C. The pellet was resuspended in BRB80 supplemented with 20 μM taxol.

### ADP treatment with hexokinase

The ADP solution was treated with hexokinase (Sigma-Aldrich, St. Louis, MO, USA) and D-glucose to remove contaminating ATP as described (You *et al*, 2017). A 50 mM ADP solution was incubated with 200 mM D-glucose, 2 mM MgClL, and 0.04 U/μl hexokinase at room temperature for 2 hours. After incubation, the enzyme was removed from the solution using an Amicon Ultra 0.5 filter (Merck). The solution was then aliquoted and stored at -20°C.

### TIRF single-molecule motility assays

TIRF assays using porcine microtubules were performed as described (Kita & Niwa, 2024). Glass chambers were prepared by acid washing as previously described (Kita & Niwa, 2024). Glass chambers were coated with PLL-PEG-biotin (SuSoS, Dübendorf, Switzerland) and streptavidin (Wako, Osaka, Japan). Polymerized microtubules were flowed into flow chambers and allowed to adhere for 5–10 min. Unbound microtubules were washed away using assay buffer (90 mM HEPES-KOH pH 7.4, 50 mM KCH_3_COO, 2 mM Mg(CH_3_COO)_2_, 1 mM EGTA, 10% glycerol, 0.1 mg/ml biotin–BSA, 0.2 mg/ml kappa-casein, 0.5% Pluronic F127, specified nucleotides with Mg diluted to indicated concentrations, and an oxygen scavenging system composed of PCA/PCD/Trolox). Purified KIF1A and KLP-6 were diluted to final concentrations of 10–20 pM and 100–400 pM, respectively, in the assay buffer. Then, the solution was flowed into the glass chamber. An ECLIPSE Ti2-E microscope equipped with a CFI Apochromat TIRF 100XC Oil objective lens (1.49 NA), an Andor iXion life 897 camera and a Ti2-LAPP illumination system (Nikon, Tokyo, Japan) was used to observe the motility. NIS-Elements AR software ver. 5.2 (Nikon) was used to control the system. All observations were completed before the motor slowed down due to ATP reduction. All analyses were conducted using kymographs generated by ImageJ software (Schneider *et al*, 2012). Unidirectional lines greater than 3 pixels along x axis (corresponding to approximately 318 nm) were counted as processive runs and the other signals were not counted. To quantify the maximum ATPase activity, segmented velocity was measured, with pause periods excluded. Nonlinear least-squares curve fitting was performed using python 3.0.

### Structural modeling

Simulation models of the KLP-6 motor domain in complex with a tubulin heterodimer were computationally constructed for both the ATP-bound and ADP-bound states, as no such structures have been reported. Cryo-EM structures of KIF1A (ATP- and microtubule-bound: PDB ID 8UTN; ADP- and microtubule-bound: PDB ID 8UTR) (Benoit *et al*, 2024) were used as templates due to the high structural conservation between the motor domains of KLP-6 and KIF1A, except for Loop-3 (Fig. S3B). To account for this difference, Loop-3 in both template structures was replaced with the corresponding loop from the autoinhibited full-length structure of KLP-6 (PDB ID 7WRG) (Wang *et al*, 2022) using PyMOL version 2.5. Based on the resulting chimeric structures, homology modeling was performed using MODELLER version 10.4 (Šali & Blundell, 1993) to generate microtubule-bound KLP-6 models for each nucleotide state. A total of 100 models were generated for each nucleotide bound state structure and top-scoring model was selected for MD simulations with discrete optimized protein energy score (Shen & Sali, 2006).

### MD simulations

Parameter and topology files for the complexes were generated using CHARMM-GUI (Jo *et al*, 2008) with the CHARMM36m force field (Huang *et al*, 2017). The MODELLER-generated structures were solvated in a cubic box of TIP3P water molecules, extending 10 Å beyond the solute in all directions. Sodium ions (NaL) were added to neutralize the system, followed by the addition of sodium and chloride (ClL) ions to achieve a physiological ionic strength of 150 mM. Energy minimization and molecular dynamics simulations were performed using GROMACS version 2023.2. The system underwent 200 steps of steepest descent minimization with positional restraints of 10 kcal/mol/Å² applied to heavy atoms of proteins and nucleotides, followed by an additional 200 steps without restraints. The system was then gradually heated to 300 K over 200 ps under constant volume (NVT) conditions with periodic boundary conditions (PBC), while maintaining positional restraints of 10 kcal/mol/Å² on heavy atoms. Initial velocities were assigned randomly for each simulation. Subsequently, constant temperature (300 K) and constant pressure (1 bar) equilibration (NPT ensemble) was performed in eight stages. The first stage consisted of a 100 ps NPT equilibration with 10 kcal/mol/Å² restraints on heavy atoms. In the following six stages, the restraints were gradually reduced to 5, 2, 1, 0.5, 0.2, and 0.1 kcal/mol/Å² in 100 ps intervals. The final equilibration step was conducted without any restraints for 100 ps. Production runs were carried out for 100 ns in the NPT ensemble at 300 K and 1 atm with PBC. Temperature and pressure were maintained using the V-rescale thermostat (Bussi *et al*, 2007) and the C-rescale barostat (Bernetti & Bussi, 2020), respectively. Long-range electrostatic interactions were calculated using the particle-mesh Ewald method (Essmann *et al*, 1995), and a 10 Å cutoff was applied to truncate non-bonded interactions. All simulations used a 2 fs time step, and hydrogen atoms were constrained using the LINCS algorithm (Hess, 2008). Ten independent 100 ns simulations were performed for each construct.

### Analysis for MD simulations

Trajectories were analyzed every 0.1 ns over the final 80 ns using CPPTRAJ (Roe & Cheatham, 2013), unless otherwise specified. Root-mean-square fluctuations (RMSFs) of the nucleotides were calculated relative to the average structure obtained from the final 80 ns of each trajectory, or up to the point at which the nucleotide dissociated from the protein (in our simulations, ADP fully dissociated from the motor domain in one out of ten runs). Secondary structure analysis was conducted using the DSSP algorithm (Kabsch & Sander, 1983). The probability of the α2a helix (residues 113–117) maintaining its helical conformation was calculated for each trajectory. The average number of hydrogen bonds formed per frame between each residue and the nucleotide was calculated across all trajectories from ten runs. Residue–residue distances between the loop-3 (K66) and P-loop (G112) were calculated across all trajectories from ten runs. Average residue–residue distances between the α2a helix and surrounding residues were measured over the final 80 ns of the trajectories across all runs, except for runs 2 and 7 of the ADP-bound WT KLP-6 simulations, in which the α2a helix was completely disrupted for more than 10 ns. For run 2, the 20–65 ns interval prior to helix disruption was used for analysis; run 7 was excluded entirely (see Fig. S6). Solvent-accessible surface area (SASA) analysis was conducted using the LCPO algorithm (Weiser *et al*, 1999). The same trajectory ranges used in the residue–residue distance analysis between the α2a helix and surrounding residues were applied here, but the average SASA was calculated separately for each run.

### Statistical Analyses and Graph Preparation

Statistical analyses were performed using Graph Pad Prism version 9. Statistical methods are described in the figure legends. Amino acid sequences of kinesins are aligned using CLUSTALW on ESPript 3 (Robert & Gouet, 2014). Structural images were generated using PyMOL version 2.5 and exported in PNG format. Graphs were created with python 3.0 and Graph Pad Prism version 9, exported in PNG format, and aligned using Adobe Illustrator 2023.

## Acknowledgments

We would like to thank the members of Niwa lab for useful discussions. T.K. was supported by JSPS KAKENHI (grant no. JP23KJ0168). SN was supported by JSPS KAKENHI (grant no. JP23H02472).

## Author contributions

T.K. writing–original draft; S.N. writing–review & editing; T.K. visualization; T.K. investigation; S.N. conceptualization; S.N. supervision; T.K. and S.N. funding acquisition.

## Statement

During the preparation of this work the authors used ChatGPT4o in order to check English grammar and improve English writing. After using this tool, the authors reviewed and edited the content as needed and take full responsibility for the content of the publication.

---

**Movie 1 Representative MD simulation of the WT KLP-6 motor domain in the ATP-bound state.**

Secondary structure elements are colored as follows: α2a helix (indigo) and P-loop (light blue). α- and β-tubulin are shown in green. The α2a helix maintains a stable helical conformation, and ATP remains stably associated with the P-loop.

**Movie 2 Representative MD simulation of the WT KLP-6 motor domain in the ADP-bound state.**

The color scheme is identical to that used in Movie 1. The helical conformation of the α2a helix is disrupted, and ADP binding is unstable.

**Movie 3 Representative MD simulation of the KLP-6(SIV/TMM) motor domain in the ATP-bound state.**

The color scheme is identical to that used in Movie 1. The α2a helix maintains a stable helical conformation, and ATP remains stably associated with the P-loop.

**Movie 4 Representative MD simulation of the KLP-6(SIV/TMM) motor domain in the ADP-bound state.**

The color scheme is identical to that used in Movie 1. The α2a helix maintains a stable helical conformation, and ADP remains stably associated with the P-loop.

