## Supplemental Information for "KLP-6 is a kinesin superfamily protein resistant to ADP inhibition"

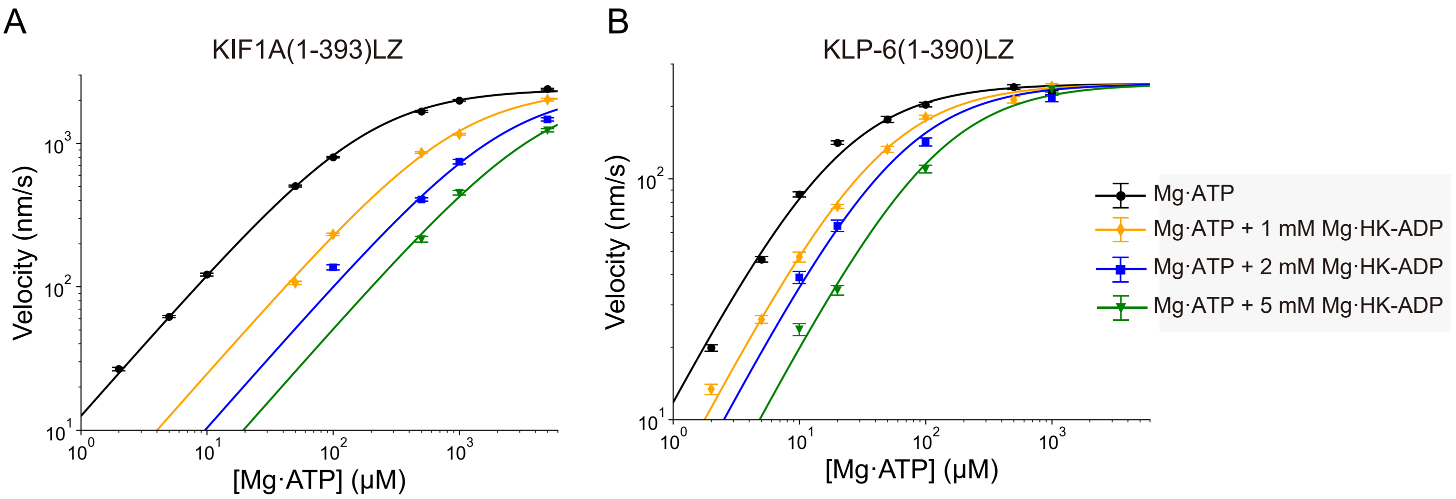


C

| ADP | KIF1A(1-393)LZ | KLP-6(1-390)LZ |
|---|---|---|
| 1 mM | 247 ± 3 μM | 901 ± 31 μM |
| 2 mM | 180 ± 4 μM | 1000 ± 69 μM |
| 5 mM | 214 ± 6 μM | 1051 ± 44 μM |

Kita and Niwa Figure S1

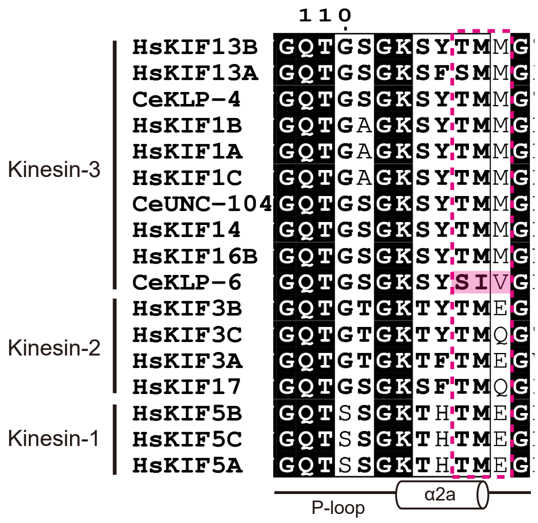


Kita and Niwa Figure S2

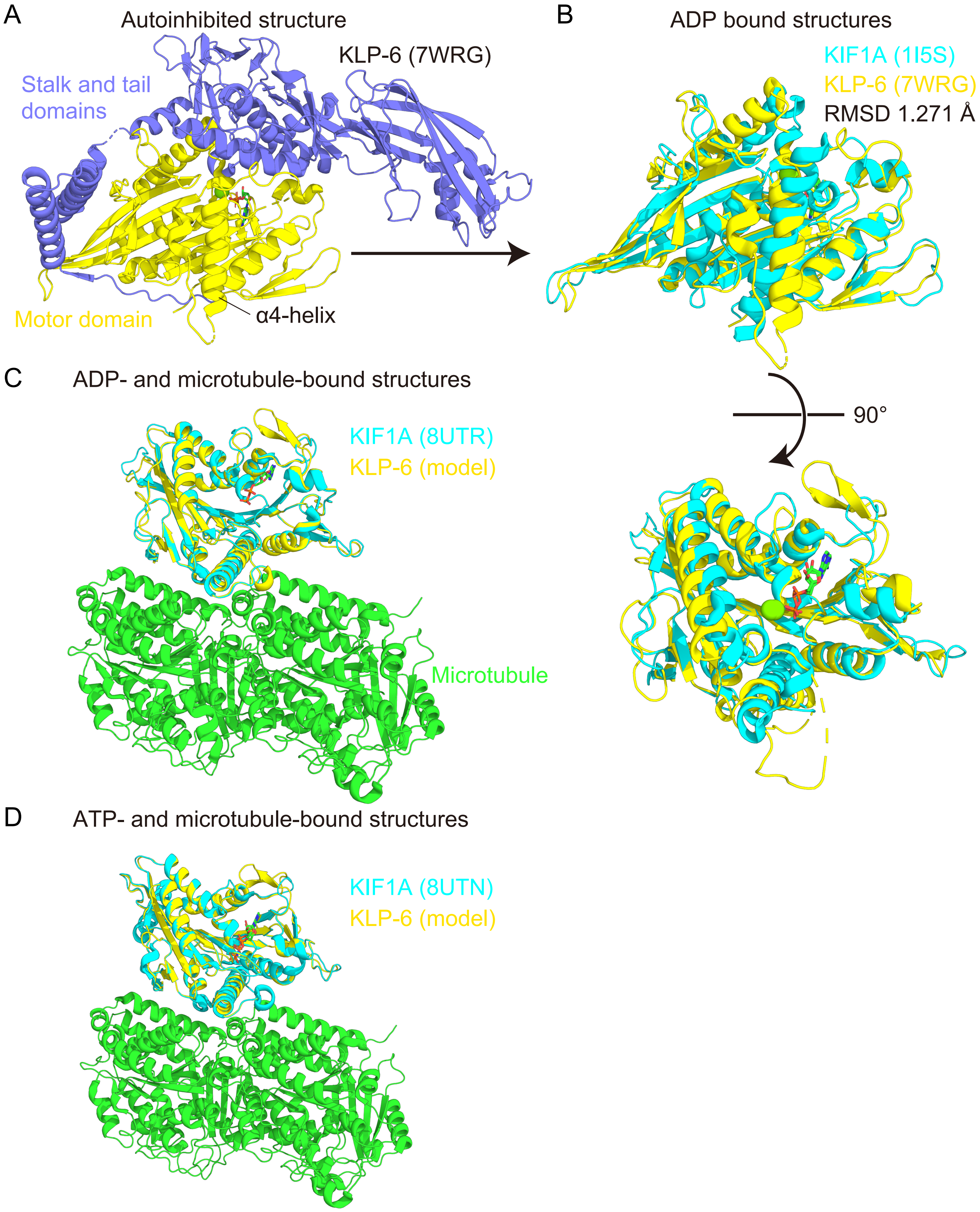


Kita and Niwa Figure S3

A

ATP-bound state

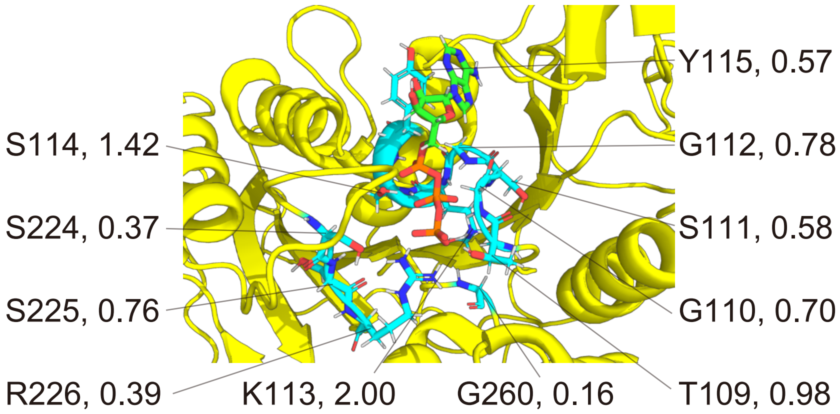


B

ADP-bound state

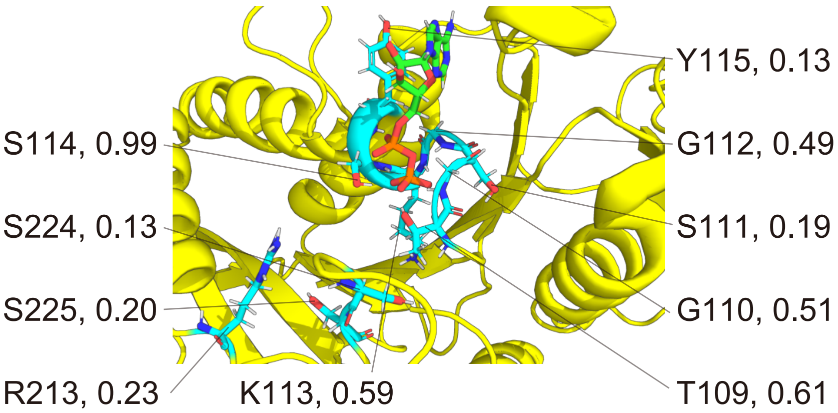


Kita and Niwa Figure S4

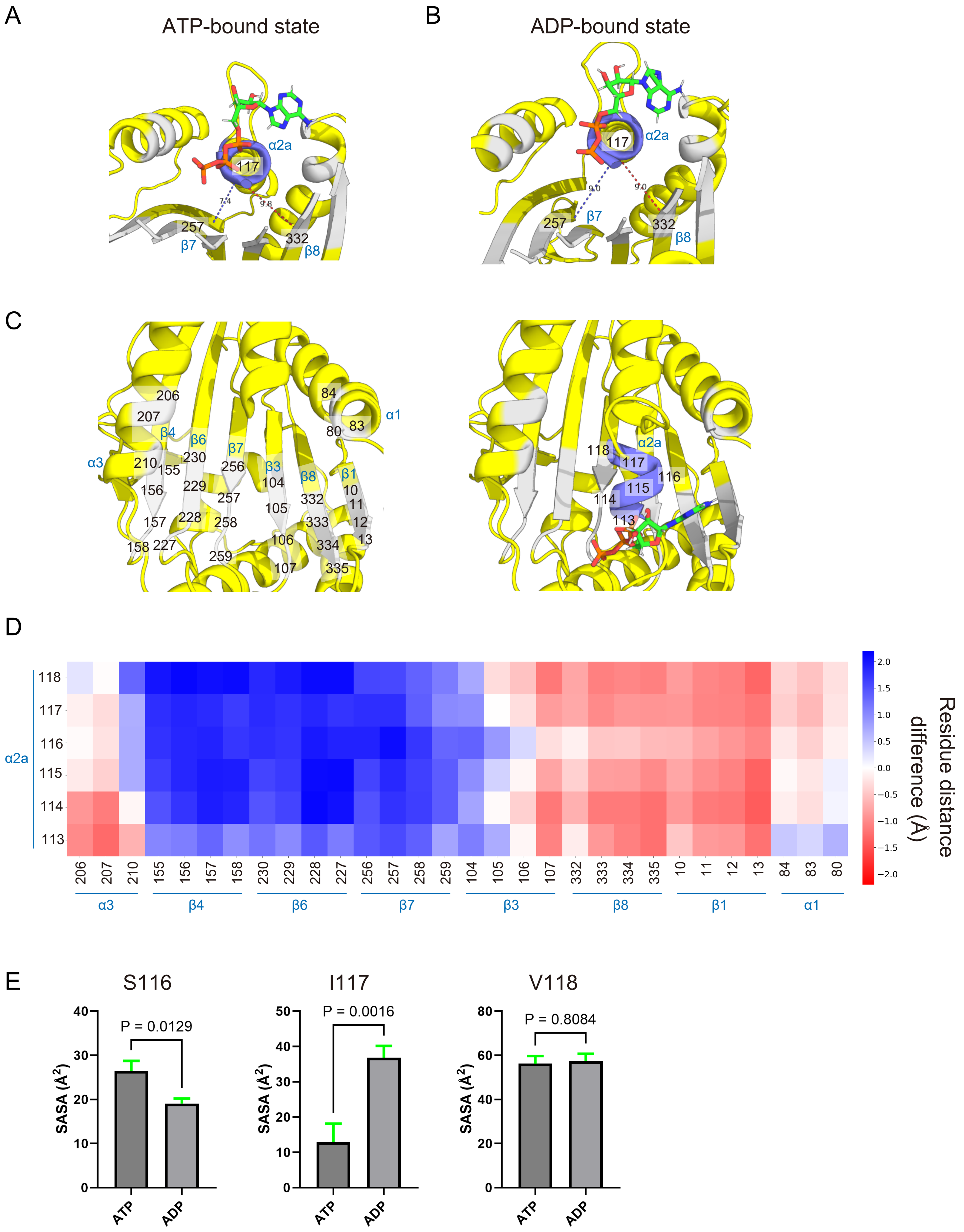


Kita and Niwa Figure S5

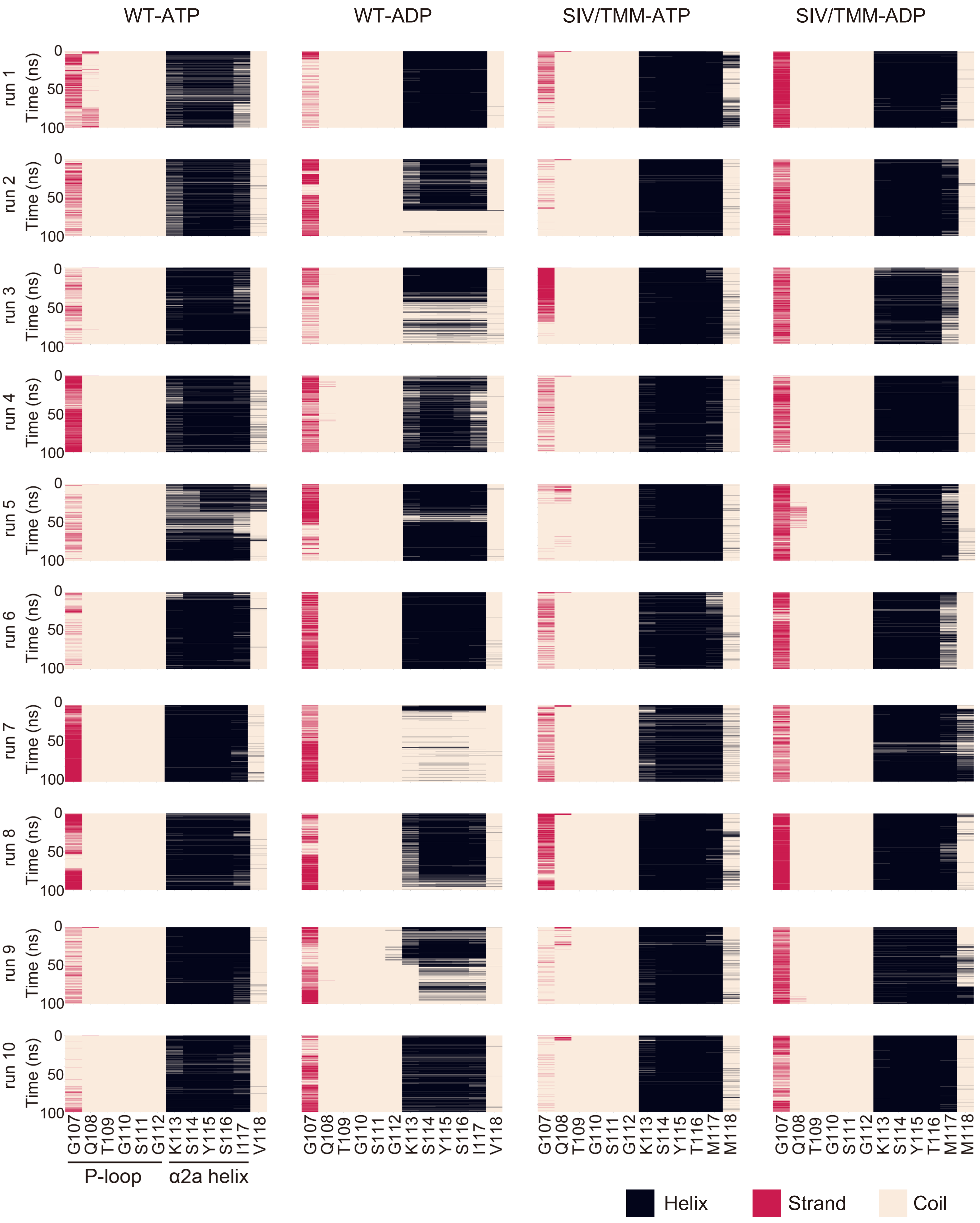


Kita and Niwa Figure S6

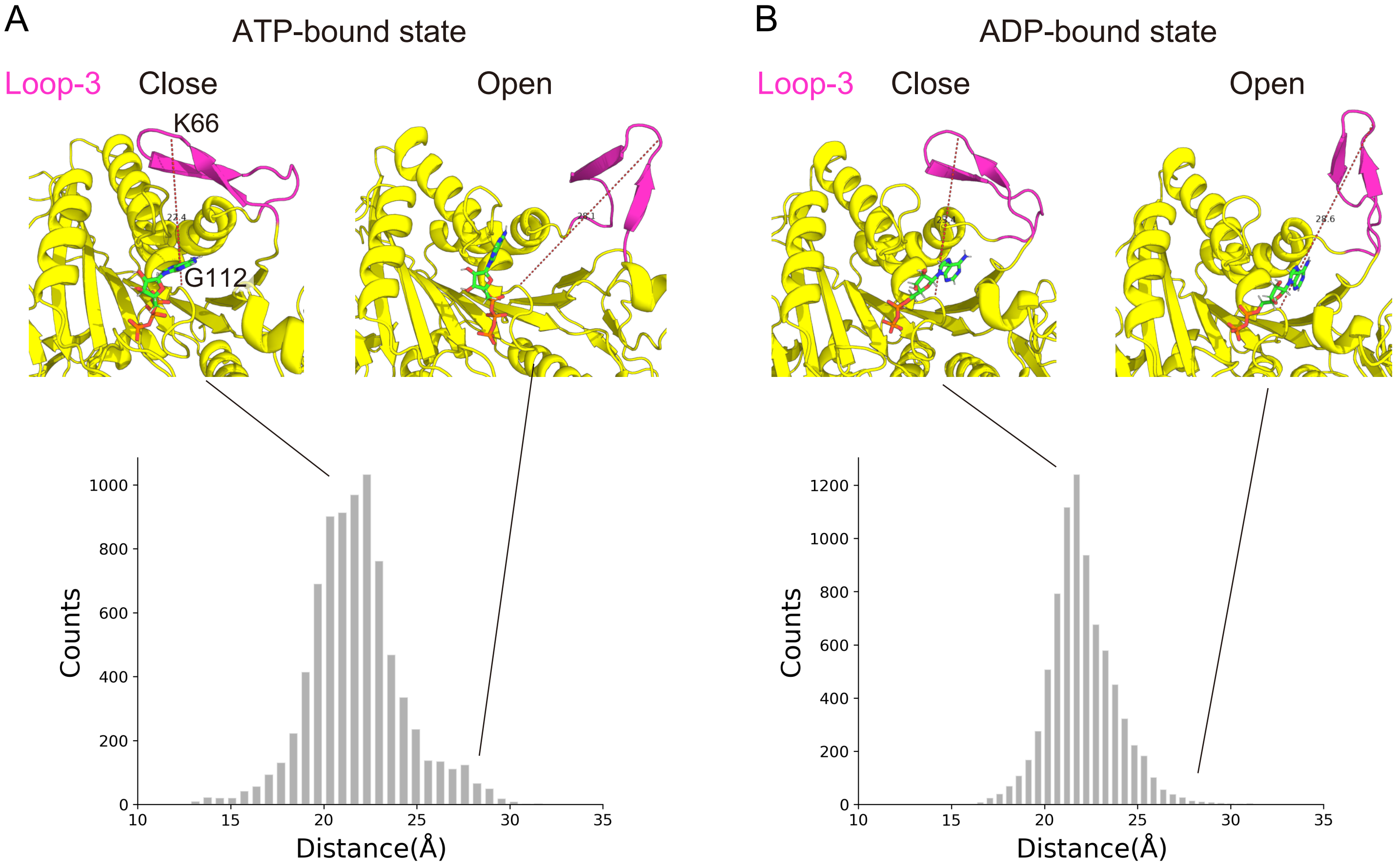


Kita and Niwa Figure S7

**Figure S1 Inhibition constant measured at different ADP concentrations.**
(A) Single-molecule velocity of KIF1A(1-393)LZ versus [Mg·ATP] in the presence of 0 mM (black circles), 1 mM (orange diamonds), 2 mM (blue square), and 5 mM (green triangle) HK-ADP. The solid lines represent the regression curves fitted to the Michaelis-Menten equation and the equation with product inhibition, respectively. Each data point represents the mean ± S.E. At least two independent experiments were conducted for each condition, with more than 96 motile particles analyzed. The data for 0 mM and 1 mM HK-ADP are identical to those shown in Fig. 2B.
(B) Single-molecule velocity of KLP-6(1-390)LZ versus [Mg·ATP] in the presence of 0 mM (black circles), 1 mM (orange diamonds), 2 mM (blue square), and 5 mM (green triangle) HK-ADP. The solid lines represent the regression curves fitted to the Michaelis-Menten equation and the equation with product inhibition, respectively. Each data point represents the mean ± S.E. At least two independent experiments were conducted for each condition, with more than 103 motile particles analyzed. The data for 0 mM and 1 mM HK-ADP are identical to those shown in Fig. 2B.
(C) Table of inhibition constants $K_I$ determined at varying ADP concentrations.

**Figure S2 Sequence comparisons of the region around the P-loop and $\alpha$2a helix.**
KLP-6-specific residues (SIV) are highlighted in magenta. In this region, the first threonine and second methionine are also conserved among kinesin-1 and kinesin-2 family members.

**Figure S3 Structural modeling for MD simulations.**
(A) Structure of autoinhibited full-length KLP-6 (PDB ID: 7WRG). The motor domain is shown in yellow, while the stalk and tail domains are colored in indigo. In this autoinhibited conformation, extension of the α4 helix is disrupted by the interaction with the stalk and tail domains.
(B) Structures of the motor domains of KLP-6 (PDB ID: 7WRG) and KIF1A (PDB ID: 1I5S). The KLP-6 motor domain was extracted from the full-length structure in (A) by removing all domains except the motor domain for visualization. The two motor domains were well aligned using PyMOL (RMSD: 1.271 Å), with the exception of Loop-3 and the α4-helix. The difference in the α4 helix conformation is attributed to the presence of the autoinhibitory stalk and tail domains in the KLP-6 structure shown in (A).
(C) Structural model of the KLP-6 motor domain bound to ADP and a tubulin heterodimer (shown in yellow). This model was generated by homology modeling based on the KIF1A structure (PDB ID: 8UTR, shown in light blue)
(D) Structural model of the KLP-6 motor domain bound to ATP and a tubulin heterodimer (shown in yellow). This model was generated by homology modeling based on the KIF1A structure (PDB ID: 8UTN, shown in light blue)

**Figure S4 Residues forming hydrogen bonds with the nucleotide in KLP-6.**

(A) Residues forming hydrogen bonds with ATP in KLP-6, as identified from MD simulations. The average number of hydrogen bonds per frame is indicated for each residue. Residues that form more than 0.1 hydrogen bonds per frame on average are listed and highlighted in light blue.

(B) Residues forming hydrogen bonds with ADP in KLP-6, as identified from MD simulations. The average number of hydrogen bonds per frame is indicated for each residue. Residues that form more than 0.1 hydrogen bonds per frame on average are listed and highlighted in light blue.

**Figure S5 Conformational differences of the α2a helix in ATP- and ADP-bound states of KLP-6.**

(A) Representative conformation of the α2a helix in the ATP-bound state of KLP-6 obtained from MD simulation. The α2a helix and surrounding residues are colored in indigo and gray, respectively. Distances from residue 117 in the α2a helix to residue 257 in the β7 sheet and residue 332 in the β8 sheet are shown.

(B) Representative conformation of the α2a helix in the ADP-bound state obtained from MD simulation. The color scheme is the same as in (A). Distances from residue 117 in the α2a helix to residue 257 in the β7 sheet and residue 332 in the β8 sheet are shown.

(C) Residues for which distances to the α2a helix were measured. The color scheme is the same as in (A).

(D) Differences in residue–residue distances between the ATP-bound and ADP-bound states of KLP-6, as determined from MD simulations. The x- and y-axes represent residues in the α2a helix and its surrounding regions, respectively, with secondary structure elements indicated. Residue pairs that are closer in the ATP-bound state are shown in blue, while those closer in the ADP-bound state are shown in red. The magnitude of the distance change is reflected by color intensity.

(E) Solvent-accessible surface area (SASA) of S116, I117, and V118 in the ATP- and ADP-bound states of KLP-6, as determined from MD simulations. Bars and error bars represent the mean and SEM, respectively. Statistical significance was assessed using Student's t-test.

**Figure S6 Secondary structure analysis of ATP-bound KLP-6 (WT-ATP), ADP-bound KLP-6 (WT-ADP), ATP-bound KLP-6(SIV/TMM) (SIV/TMM-ATP), and ADP-bound KLP-6(SIV/TMM) (SIV/TMM-ADP).**

The analysis was performed across ten independent simulations for each construct. Each graph shows the time-dependent changes in secondary structure for residues near the P-loop and the α2a helix. Black, red, and pale orange indicate helix (including α-helix, $3_{10}$-helix, or $\pi$-helix), strand (including residues in isolated β-bridges or extended strands that participate in β-sheets), and coil (including hydrogen-bonded turns, bends, loops, or irregular structures), respectively.

**Figure S7. Conformation of loop-3 in the ATP- and ADP-bound states of KLP-6.**

(A) Histogram showing the distance between K66 in loop-3 and G112 in the P-loop in the ATP-bound state of KLP-6, as determined from MD simulations. Representative structures of loop-3 in closed and open conformations are shown above. Loop-3 is colored in magenta, and the measured distance is indicated.

(B) Histogram showing the distance between K66 in loop-3 and G112 in the P-loop in the ADP-bound state of KLP-6, as determined from MD simulations. Representative structures of loop-3 in closed and open conformations are shown above. Loop-3 is colored in magenta, and the measured distance is indicated.
